# Re-shaping the immune response to influenza vaccination in a host with immune memory from influenza

**DOI:** 10.1101/2025.10.01.678836

**Authors:** Chantelle L. White, Lara Mengu, Siva K. Gandhapudi, Katherine A. Richards, Andrea J. Sant

**Affiliations:** David H. Smith Center for Vaccine Biology and Immunology, Department of Microbiology and Immunology, University of Rochester, Rochester, NY 14642; Department of Microbiology, Immunology and Molecular Genetics, University of Kentucky School of Medicine, Lexington, KY 40536

## Abstract

Although CD4 T cells are critical orchestrators of protective immunity to viral respiratory pathogens, vaccine strategies that optimize generation of these cells have yet to be prioritized. In this manuscript, to mimic the typical human vaccine recipient using a mouse model, we evaluated the impact of previous influenza infection on the adaptive immune response elicited by the recombinant influenza vaccine Flublok, co-delivered with either AddaVax, an MF59 mimetic or a nanolipoparticle innate activator R-DOTAP. In the context of influenza B infection memory, a repolarization of the responding CD4 T cells and dramatic change in the fate of the vaccine-elicited CD4 T cells was discovered. A rapidly evolving CD4 T cell response enriched in TNF-α and IFN-γ was observed, with the HA-B-specific CD4 T cells also displaying increased expression of chemokine receptors associated with lung homing potential and ultimate accumulation in the lung tissue. Unexpectedly, similar shifts in the features of the H3-specific CD4 T cell and antibody response were also observed, drawn from the naïve repertoire. These results are consistent with the view the microenvironment of the vaccine draining lymph node, developed in the context of immune memory, rather than infection-induced CD4 T cell imprinting, plays the decisive role in the functional phenotype, magnitude, and fate of vaccine-elicited CD4 T cells. These results have important implications for both pre-clinical models of vaccination and future vaccine strategies.

## Introduction

Most current vaccine strategies to protect humans from respiratory infections focus on elicitation of antibodies that block infection such as hemagglutinin or Spike for protection from influenza or SARS-CoV-2, respectively. The challenge of these approaches is the emergence of variant strains that evade this antibody-mediated protective immunity by mutation and selection *in vivo*, creating drift variants that require vaccine reformulation, composed of the new drifted strains. Because of this limitation, there is a growing focus on vaccine strategies that enhance protective immunity by augmenting cellular immunity to complement antibody-mediated immunity ^1–4^. T cell mediated immunity is notable in that it can target genetically conserved pathogen-derived epitopes that persist in drifted variants. Additionally, CD4 T cells are particularly important because of the diverse effector functions they convey, including production of cytokines such as IL-4 that promotes B cell responses, IL-2 that fosters CD8 T cell expansion and memory, and IFN-γ that has direct anti-viral effects and that can upregulate proteins associated with antigen processing and presentation ^5–8^. Current efforts to leverage cellular immunity in vaccines include peptide-based vaccines ^9,10^, attenuated viruses, and formulation of protein-based vaccines with immunomodulatory compounds and innate activators that promote cellular immunity ^11–13^. As part of these efforts, we have examined the capabilities of adjuvants that have been used in human vaccines to potentiate CD4 T cell immunity. Our studies revealed that the co-administration of Flublok, a licensed recombinant protein based influenza vaccine ^14,15^ with the novel innate activator R-DOTAP (“R-DP”), outperforms the MF59 mimetic Addavax (AdVx), in the generation of robust, CD4 T cell populations with multifunctional phenotypes, even when combined with the potent TLR agonist CpG.

Despite these promising results, when designing translationally relevant, preclinical models to address vaccine efficacy, it is important to account for the fact that most of the individuals who receive these vaccines will rarely be immunologically naïve to influenza. By 2 years of age, most children will have had their first encounter with influenza through infection ^16^. The events that occur upon vaccination of a non-naïve host have not been explored mechanistically and will likely depend on factors such as the influenza vaccine formulation and whether any innate activator is used (reviewed in ^11,17–19^). The recruitment of infection-induced T cells to peripheral vaccination can be envisioned to impact the immune response in several ways, based on studies of secondary infections and prime-boost vaccine strategies. The first is that the kinetics of the responses is likely to be more rapid. The T cell receptor (TcR) signaling threshold for the reactivation of memory CD4 T cells is lower than naïve CD4 T cells ^20^, leading to possible earlier T cell detection of peptide:MHC complexes on the surface of antigen presenting cells (APC) after vaccination. Also, memory T cells will increase the precursor frequency which together with increased sensitivity to antigen would likely elicit a more rapid T cell response, with perhaps a cytokine profile altered by the dominance of the infection-induced memory cells.

Despite these suspected events, the quantitative and qualitative impact of this recruitment of memory cells to vaccine responses is not known. To address these possibilities, we have established an experimental system to examine peripheral influenza vaccine responses in animals that have influenza-specific memory initiated by infection. Mice with established memory have been vaccinated with a licensed recombinant protein vaccine, introduced with either a standard oil in water adjuvant, (Addavax, “AdVx”) or a novel enantio-specific cationic lipid nanoparticle (R-DOTAP, “R-DP”) to enhance vaccine responses, as we have previously shown ^21,22^. Our previous studies with recombinant influenza protein vaccination in naïve mice revealed that the R-DP elicited epitope specific CD4 T cells of greater abundance and complexity than AdVx with or without added TLR ligand, CpG ^23,24^.

In the studies reported here, we have addressed the impact of immune memory on vaccine responses. Upon primary vaccination of mice with a history of infection with influenza B (IBV) with either Flublok/AdVx or Flublok/R-DP, we uncovered a drastic shift in the quantity, response kinetics, effector phenotype and fate after direct comparison to responses in immunologically naïve mice, as well as their fate for both immune activator systems. Interestingly, the recruitment of naïve CD4 T cells specific the alternate HA protein (H3), for which there was no memory, displayed parallel trajectories as the HA-B specific cells, as well as the H3 specific antibody response, suggesting that the vaccine draining lymph node (dLN) where these responses to vaccination take place reshape the response phenotype and kinetics of both memory and naïve cells.

## Materials and Methods

### Mice and Ethics Statement

C57BL/6 (“B6”) female mice were purchased from the National Cancer Institute (NCI, Fredrick National Laboratory, Bethesda, MD, USA). Mice were typically used between the ages of 12 -16 weeks and were housed in a specific pathogen-free facility at the University of Rochester Medical Center as required by institutional guidelines. The mouse experiments followed AAALAC International, the Animal Welfare Act, and the PHS Guide and were approved by the University of Rochester Committee on Animal Resources, Animal Welfare Assurance Number D16-00188. The protocol for the described studies was originally approved on 4 March 2006 (protocol no. 2006-030) and was evaluated and re-approved every 36 months. The most recent review and approval was on 21 December 2023.

### Influenza Virus Infections

C57BL/6 mice were anesthetized by intraperitoneal injection with tribromoethanol (Avertin, 14 μL/mg body weight) and then intranasally infected with B/Brisbane/60/2008 at 1200 PFU, adjusted to 30 μL in PBS. The mice were monitored for health and weight until their use at >30 days post infection.

### Preparation of R-DP Nanoparticles and vaccine formulations

A Flublok Quadrivalent (Sanofi) 2019–2020 formula containing recombinant hemagglutinin proteins derived from the influenza strains A/Brisbane/02/2018 (H1N1), A/Kansas/14/2017 (H3N2), B/Maryland/15/2016, and B/Phuket/3073/2013 was obtained from the manufacturer through the University of Rochester Medical Center Pharmacy. Each 0.5 mL dose of Flublok contains 45 μg of each HA (180 μg total HA) in PBS and 27.5 μg of Tween-20.

Current good manufacturing practice–grade (CGMP) R-DP was provided by PDS Biotechnology Corporation (Florham Park, NJ, USA). Flublok vaccine in PBS buffer was diluted with 280 mM sucrose for R-DP or PBS for AdVx. Before administration to animals, the vaccines were brought to room temperature. Using a pipette to form a uniform suspension, the Flublok vaccine (10 μL equal to 3.6 μg total HA, or 0.9 μg of each HA protein) was added at a 1:1 ratio with the R-DP nanoparticles (6 mg/mL in 280 mM sucrose), as previously described ^24^. For AdVx, an equal volume of Flublok in PBS and AdVax was prepared. For vaccination of all cohorts of mice, 100 μL of Flublok with or without the immune activators was used for each dose, delivered subcutaneously in 50 μL aliquots to each of the rear footpads. Mice were sampled at the sites and times post vaccination indicated in each Figure.

### Peptides

16-mer peptides were obtained from Biopeptide (San Diego, CA, USA). For the Influenza B single peptide stimulation, the immunodominant peptide HA-B p6/7: ^23^TSSNSPHVVKTATQGE^38^ (“HA-B”). For detection of H3-specific CD4 T cells, the two adjacent peptides, p35 and p36 (p35/36) (p35: ^203^TNNDQISLYAQASGRIT^219^, p36: ^209^SLYAQASGRITVSTKRS^225^), were pooled and referred to as “H3”. Single epitope stimulations were performed at a final concentration of 2 μM. These peptides have been defined previously ^23^.

### Intravascular Antibody Labeling

At the designated time post-infection or vaccination, mice were anaesthetized through inhalation of isoflurane and retro-orbitally injected with 3 μg anti-mouse CD45 (30-F11, Tonbo) in a total volume of 100 μL ^25^. After 4 minutes, mice were euthanized with an overdose of Avertin and the lungs, mediastinal lymph nodes (mLN), spleen and popliteal lymph nodes were harvested.

### Tissue Processing and CD4 T cell purification

Using a razor blade, the lungs were minced into fine segments, placed in GentleMACS tubes (Miltenyi Biotec) and incubated on a shaker for 1 hour at 37°C, in lung digestion media (collagenase type II (10^4^ units/ml) and DNase I (30 μg/ml)) in RPMI (Gibco) supplemented with 2.5% fetal bovine serum (FBS, Gibco) and 10 mM HEPES (Gibco). After incubation, GentleMACS tubes were connected to the GentleMACS Dissociator. Samples were then passed through a 40-μM sterile nylon mesh. The mLN, spleen and pLN were mechanically disrupted using the back of a 3 mL syringe and passed through 40-μM sterile nylon mesh. All samples were washed with Dulbecco’s modified Eagle medium (DMEM, Gibco) supplemented with 1% gentamicin and 10% heat-inactivated FBS and then red blood cell depleted using ACK lysis buffer (0.15 M NH_4_Cl, 1.0 mM KHCO_3_, 0.1 mM NaEDTA, pH 7.2). CD4+ T cells were enriched by negative selection via MACS (Miltenyi Biotec) using the manufacturer’s protocol. Purified CD4+ T cells were treated to surface and intracellular staining and evaluated by spectral flow cytometry, as described below.

### Flow cytometry

To evaluate the antigen-specific response, CD4 T cell enriched populations were resuspended at 300,000 cells per well in media (complete DMEM media with 10% FBS (Fisher Scientific, Gibco)), and co-cultured with splenocytes at 500,000 cells per well, from naïve, age and sex matched mice, in sterile U-bottom plates. The antigen-specific responses were determined by stimulating the cells with the HA-B and H3 peptides, described above, at 2 μM per well. Plates were incubated at 37^◦^C and 5% CO_2_. After one hour of incubation, a combination of two protein-trafficking inhibitors, monensin (GolgiStop, BD Biosciences) and brefeldin (GolgiPlug, BD Biosciences), were added to the culture. At this time, an anti-CD107a antibody (clone 1D4B, Invitrogen) was also added to the culture at a 1:200 dilution. Plates were incubated for an additional 9 h at 37^◦^C and 5% CO_2_ and then transferred to 4^◦^C until staining. After stimulation, samples were transferred to V-bottom tissue culture plates and washed twice with PBS. Cells were stained for viability with LIVE/DEAD Fixable Blue Dead Cell Stain (Fisher Scientific, Invitrogen) for 30 min at 4 ◦C. Next, the cells were washed twice with FC stain buffer (PBS, 2% heat-inactivated FBS, 0.01% NaN_3_) and then resuspended in anti-CD16/CD32 (FC Block, clone 2.4G2, BD Biosciences) for 15 min at 4^◦^C. Without washing, the cells were stained for 30 min at 4^◦^C with a surface stain master mix **Supplemental Table 1**. For the intracellular staining, cells were washed twice with FC stain buffer and then fixed and permeabilized using FoxP3 Fixation/Permeabilization buffer (Fisher Scientific, eBiosciences) and placed at 4^◦^C for 1 h. After washing the cells twice with FoxP3 wash buffer (eBiosciences), the cells were incubated in FC Block and an intracellular stain master mix (**Supplemental Table 1**). The cells were washed twice with FoxP3 wash buffer and then fixed in 1% paraformaldehyde for 10 min at 4^◦^C. Once fixed, the cells were washed and resuspended in FC stain buffer for data acquisition.

### Data Acquisition and Analysis

Samples were run on a 5-laser, Cytek Aurora (Cytek, Bethesda, MD, USA). Data were analyzed using FlowJo software, version 10.9.0 (Becton, Dickinson and Company, Ashland, OR, USA). A statistical analysis was conducted using GraphPad Prism software, version 10. Specific statistical tests used for each data set are indicated in the Figure Legends. p values are indicated as an asterisk using the following criteria: p > 0.05, *: p ≤ 0.05, **: p ≤ 0.01, ***: p ≤ 0.001, and ****: p ≤ 0.0001.

### ELISA

HA-specific IgG antibody responses were quantified by ELISA. High-binding plates (Corning Costar, Tewksbury, MA, USA) were coated with 200 ng/well of purified recombinant HA protein from either Influenza A/Kansas/14/2017 or Influenza B/Brisbane/60/2008. After overnight coating at 4°C, plates were washed with PBS and blocked with 3% BSA in PBS at room temperature for 1 h. Diluted serum samples (in 0.5% BSA–PBS) were added and incubated for 2 hours at RT. Plates were then washed and incubated with alkaline phosphatase–conjugated goat anti-mouse IgG (SouthernBiotech, Birmingham, AL, USA), followed by p-nitrophenyl phosphate (pNPP) substrate (Sigma, Burlington, MA, USA). Absorbance was measured at 405 nm to quantify antibody binding.

## Results

### The primary response to Flublok vaccination in naïve mice administered with two distinct immune activator systems

The primary response to vaccination in naïve mice at the peak of the response was studied first, in order to understand the baseline from which the recall would take place. **Figure 1A** shows the results from epitope-specific intracellular cytokine staining with multiparameter flow cytometry for Flublok tested for HA-B specific responses (**Figure 1A top**) and H3-specific CD4 T cell responses (**Figure 1A, bottom)**. The response magnitude is depicted for different cytokines elicited with no adjuvant (in green), which is barely detectable, and the AdVx and R-DP formulations of Flublok in blue and orange, respectively. These experiments show that the two immunomodulatory systems are distinct in their impact on Flublok responses. AdVx elicits a Th2-biased CD4 T cell response, enriched for IL-4, with a smaller population of CD4 T cells producing IL-2 and TNF-α, while the R-DP-formulated Flublok elicits a much more robust response (approximately 4-fold), enriched for IFN-γ and TNF-α, with negligible production of IL-4. Similar patterns of responses were seen for H3-specific CD4 T cells (**Figure 1A, bottom)**, although the magnitude of the response to H3 is noticeably lower than that to HA-B, likely due at least in part to the presence of both HA-B lineage proteins in the Flublok, which would double the dose. It should be noted that in this strain of mice, there are negligible CD4 T cell responses to H1 ^26^, and so this specificity has not been tracked in these studies.

**Figure 1.**
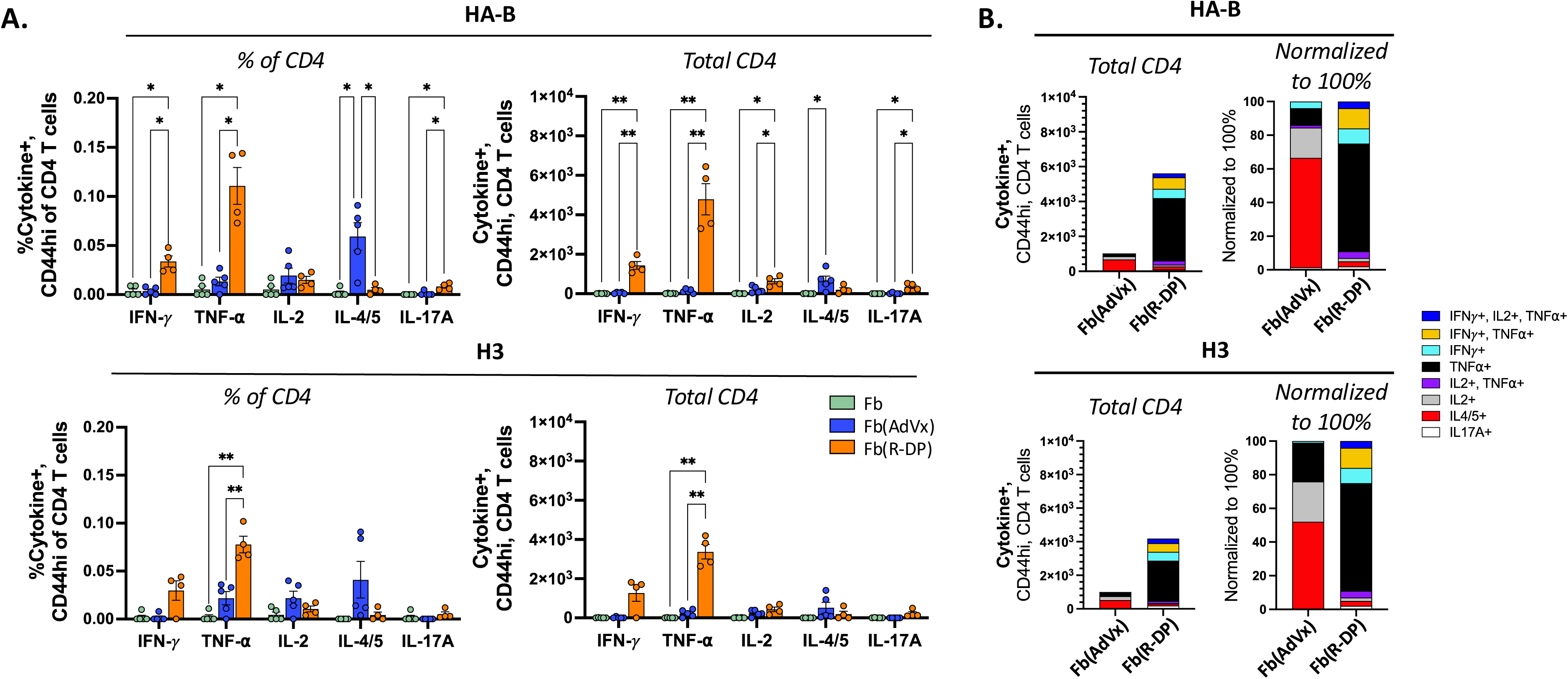
Phenotypic characterization of the primary response to Flublok formulated with the two immunomodulatory systems. Mice were immunized subcutaneously in the footpad with either Flublok alone (Fb) or Flublok combined with AddaVax (AdVx) or R-DOTAP (R-DP) and at 9 days post-vaccination, purified CD4 T cells were evaluated by ICS and flow cytometry. Panel A. IFN-*γ*, TNF-⍺, IL-2, IL-4/5, IL-17A, CD44hi, CD4 T cell populations detected upon stimulation with HA-B (top) or H3 (bottom) peptides after vaccination with Flublok alone (green), Flublok(AdVx) (blue) or Flublok(R-DP) (orange). Antigen-specific responses are quantified here as a % of CD4 T cells (left) or total per mouse (right). Bars reflect the mean with SEM. Two-way ANOVA with Geisser-Greenhouse and Fisher LSD test was used to evaluate significance. Panel B. HA-B- or H3-specific epitope-reactive cells were quantified by ICS and flow cytometry. Polyfunctionality was determined by Boolean combination gating identifying 8 distinct populations (colors indicated at right) that produced cytokine at least 2-fold above background. The height of the individual stack in the bar reflects the mean of the total cytokine producing, antigen-reactive response and the asterisk indicate the significance as follows *p < 0.05, **p < 0.01, ***p < 0.001 and ****p < 0.0001.

Use of multiparameter flow cytometry with Boolean gating was then used to assess CD4 T cells producing two or more cytokines for the vaccinated mice. These data are illustrated as stacked bar graphs in **Figure 1B**. R-DP/Flublok elicited many subsets of CD4 T cells, including IFNγ/TNFα double producers, and IFNγ/IL2/TNFα triple producers, as well as cells producing single cytokines, primarily TNF-α alone, with only minor populations of cells producing IL-4 or IL-17. In contrast, AdVx/Flubok primarily elicited CD4 T cells producing IL-4 alone with smaller populations of cells producing IL-2 or IL-17 alone or multiple cytokines. The patterns for HA-B and H3-specific CD4 T cells were similar. The distinctive patterns with alternate immune activator systems provided an excellent opportunity to analyze the response to vaccination in mice with established memory from infection.

### Influenza infection memory alters the magnitude and kinetics of the CD4 T cell response to vaccination

To examine the impact of a history of influenza infection on vaccine responses, we established immune memory via infection with influenza B virus (IBV), which contained abundant CD4 T cell epitopes, including one dominant HA-B epitope. The localization and phenotype of infection-elicited CD4 T cells was assessed at memory, prior to vaccination, using HA-B peptide stimulation and multiparameter flow cytometry. As shown in a representative experiment in **Figure 2**, most of the cytokine-producing CD4 T cells are in the spleen, followed by the lung tissue and vasculature. Multiple subsets of CD4 T cells were identified, demonstrating functional complexity within the memory compartment. CD4 T cells within the lung vasculature and spleen were largely comprised of IFN-γ and TNF-α producers, either produced alone or together, suggesting that CD4 T cells recruited into the draining lymph node (dLN) might enrich the microenvironment with Th1-associated cytokines.

**Figure 2.**
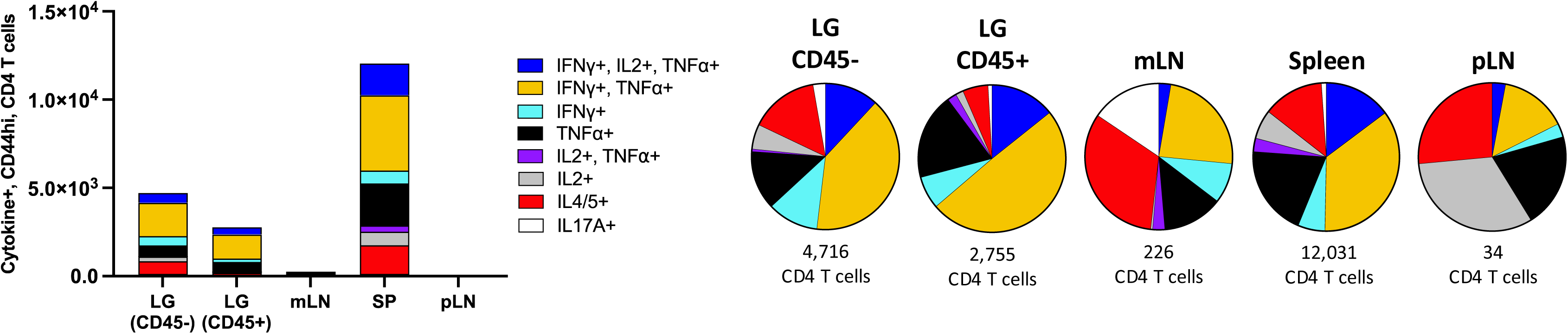
Tissue distribution and functional profile of antigen-reactive CD4 T cells at memory. C57BL/6 mice (n = 25) were infected with B/Brisbane/60/2008 and at 33 days post infection (dpi), a fluorescent CD45 antibody was retro-orbitally injected 4 minutes prior to sacrifice and in the harvested lung used to distinguish CD4 T cells localized to the lung tissue (CD45-) vs. vasculature (CD45+). Antigen-specific reactivity was determined by co-culturing purified CD4 T cells with the HA-B stimulating peptide and splenocytes from naive mice. Each unique cytokine-producing population comprising the total antigen-reactive response is individually represented here with a unique color. The height of the stacked bar graph indicates the mean total quantity of cytokine-producing CD4 T cells per mouse in each tissue. Pies reflect the mean total antigen-specific response in each tissue (total number listed below each pie), with each slice representing a unique cytokine-producing population as a fraction of the total response.

To evaluate the consequences of the infection-induced memory CD4 T cells on the elicited Flublok vaccine response, large replicate studies were initiated, where responses elicited by the AdVx/Flublok was compared with that elicited by R-DP/Flublok, using the design illustrated in **Figure 3A** in naïve or “IBV-memory” mice. In naïve mice, or in the cohorts 33 days post-infection (dpi), mice were subcutaneously vaccinated in the footpad with Flublok, formulated with either AdVx (IBV/Flublok(AdVx)) or R-DP (IBV/Flublok (R-DP)). Several key questions could be addressed with this experimental design. First, what is the functionality and abundance of CD4 T cells in the vaccinated host with IBV memory compared to the naïve animal? Second, do the unique features of two immunomodulatory systems observed in the naïve host persist in the host with memory? Finally, does the fate of the vaccine-elicited CD4 T cells differ in naïve vaccinated mice vs. those with infection memory? Because we suspected that the recruitment of memory CD4 T cells into would accelerate responses, the vaccine draining popliteal lymph node (pLN) was sampled at day 6 and day 9 post-vaccination (“dpv”), the latter time point previously determined to be peak of the response in naïve animals. CD4 T cells, purified from the pLN in both cohorts and stimulated with the dominant HA-B peptide epitope (“HA-B 6/7” or “HA-B”), were analyzed for abundance and cytokine patterns. These side-by-side analyses revealed a dramatic impact of the CD4 T cell response to vaccination when it occurs in a host with a history of infection. First, when all epitope-specific cytokine producing cells were summed (**Figure 3B, left panel**), we found that by 6 dpv, there was a sharp increase in CD4 T cell responses within influenza infection experienced, Flublok-vaccinated mice. The differences in the scales show that while responses to both vaccine formulations were enhanced in the context of immune memory, CD4 T cell responses in IBV/Flublok (R-DP) treated mice were 15-fold higher than in the IBV/Flublok (AdVx) cohort, suggesting that R-DP vaccination is more potent in recruiting or expanding infection-induced memory cells. Second, the abundance of cytokine-producing CD4 cells in the dLN over time depended on the immune history of the host (**Figure 3B right panel**). In naïve mice, antigen-specific CD4 T cells continued to accumulate from day 6 to day 9, but in the influenza-experienced mice, the responses peaked at day 6 and then declined by day 9.

**Figure 3.**
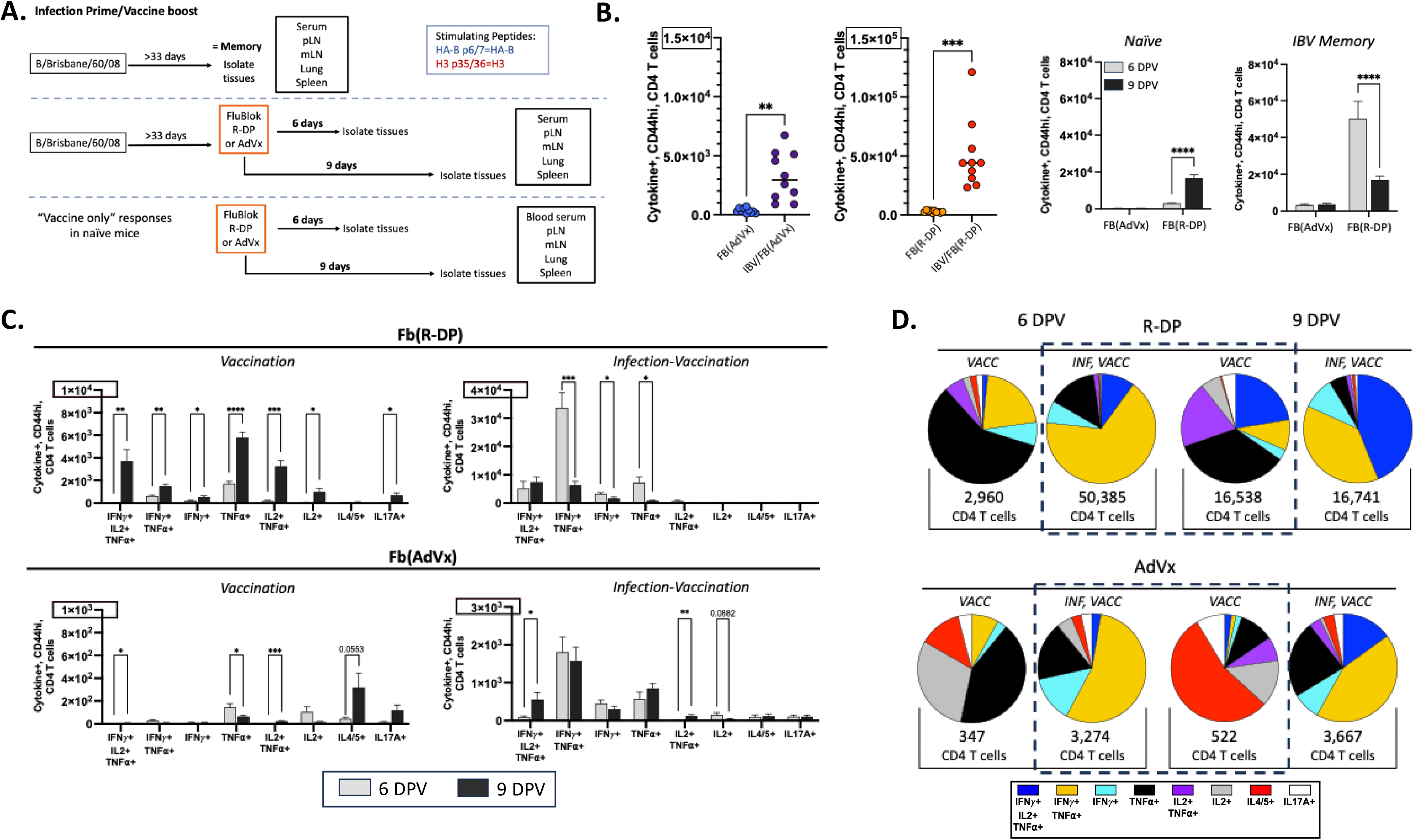
Remodeling of IBV-specific CD4 T cell responses. The experimental design to evaluate the impact of infection-induced memory on vaccine response is shown in panel A. Panel B left indicates the sum of HA-B specific cytokine-producing cells identified by ICS and flow cytometry. Each symbol represents an individual animal from 3 independent experiments with statistics performed using unpaired t test with Welches correction. B, right. The magnitude of HA-B-specific responses of naïve (left) or previously (IBV)-infected mice (right) after vaccination with Fb(AdVx) and Fb(RDP) at day 6 or day 9 post-vaccination. Bars represent with mean with SEM and a paired t test. C. Remodeling of HA-B-specific cytokine-producing CD4 T cell populations after vaccination of naïve (left) or previously infected (right) mice at 6 (grey), compared to 9 days post-vaccination (black). Bars with Flublok(R-DP) (top) or Flublok(AdVx) (bottom) reflect the mean with SEM and statistical analysis using a two-way ANOVA and Fisher’s least significant difference test. P values are indicated as follows * p < 0.05, ** p < 0.01, *** p < 0.001, and **** p < 0.0001. D. Subsets of HA-B specific CD4 T cells identified by ICS and flow cytometry from 6 to 9 days post vaccination with Flublok formulated with R-DP (top) or AdVx (bottom) shown as the mean of the total antigen-specific response at day 6 and day 9 with the total noted below each pie. Each slice, assigned a distinct color, reflects the fraction of the total response attributed to each unique cytokine-producing population.

### Reshaping of the phenotype of the vaccine elicited CD4 T cells by vaccination of host with previous infection

When subsets of HA-B specific cytokine-producing CD4 cells were quantified at day 6 and day 9, shifts in the relative abundance of CD4 T cells were observed (**Figure 3C**). In the naïve mice (left panels) the dominant cytokine-producing CD4 T cells increased from day 6 to day 9 for both R-DP (Top) and AdVx (bottom) formulated Flublok responses (note the differences in scales for R-DP and AdVx), with maximal responses boxed. In contrast, as seen in the right panels of Figure 3C, within the host with memory, instead of continued accumulation over time, many subsets of HA-B specific were diminished at day 9, particularly with Flublok/R-DP.

The overall reshaping of the CD4 T cell response to vaccination is best illustrated by pie graphs (**Figure 3D**), where the quantity of HA-B-reactive CD4 T cells was summed followed by calculation of the fraction of each of the eight distinct subsets of cytokine producing CD4 T cells. When the peak of each response is compared (shown boxed at day 9 for naïve and day 6 for IBV memory mice), both immune activator systems were remarkably reshaped by influenza memory in the host. Perhaps most striking was the response to Flublok/AdVx, where the IL-4 production (in red) was essentially extinguished and replaced by CD4 T cells producing primarily IFN-γ and TNF-α (in gold). The responses with Flublok formulated with R-DP shifted from an TNF-α dominant cytokine response (in black) with other subsets of CD4 T cells producing IL-2 and TNF-α (in purple) and those producing IL-2 and TNF-α, and IFN-γ (in blue) to a dominant population of CD4 T cells producing TNF-α and IFN-γ (in gold).

### Chemokine receptor expression on HA-specific CD4 T cells changes in a host with infection history

One intriguing possibility for the loss of subsets of cytokine-producing CD4 T cells in the dLN in the host with infection-induced memory between day 6 and day 9 is selective egress of CD4 T cells from the dLN driven by chemokine receptor expression. To evaluate this possibility, the chemokine receptor expression profiles of the HA-B epitope-specific CD4 T cells was evaluated. Th1 cells have been shown to preferentially express CXCR3, CCR5 and CXCR6 ^27–30^, allowing these cells to migrate into sites of inflammation, including the lung ^29,31^. To evaluate this, HA-B specific CD44hi, cytokine-producing CD4 T cell populations at day 6 and day 9 were assessed for their expression of CXCR3, CCR5 and CXCR6, as shown in **Figure 4A**. Sample flow plots comparing the frequency of each chemokine receptor expressing population in IBV memory mice vaccinated with Flublok (AdVx) or Flublok (R-DP) are shown **Supplemental Figure 1**. The gates identifying chemokine expressing cells were applied to IFN-γ producing populations to determine the fraction of each functional subset that was positive for each of the chemokine receptors tested. At 6 dpv (**Figure 4A**), the fraction of cytokine producing cells that expressed CXCR3, CCR5 or CXCR6 was significantly elevated in mice previously infected with IBV and then vaccinated with Flublok/R-DP (in orange). Based on the limited chemokine receptor expression in IBV/Flublok/AdVx or Flublok/R-DP vaccinated naïve mice, we conclude that neither infection history nor Flublok/R-DP vaccination is sufficient to increase the frequency of chemokine receptor expressing, antigen-reactive CD4 T cells. Instead, our data is consistent with the view that it is both the early inflammatory signals initiated by R-DP and the subsequent recruitment of memory populations into the dLN are needed for upregulation of chemokine receptors.

**Figure 4.**
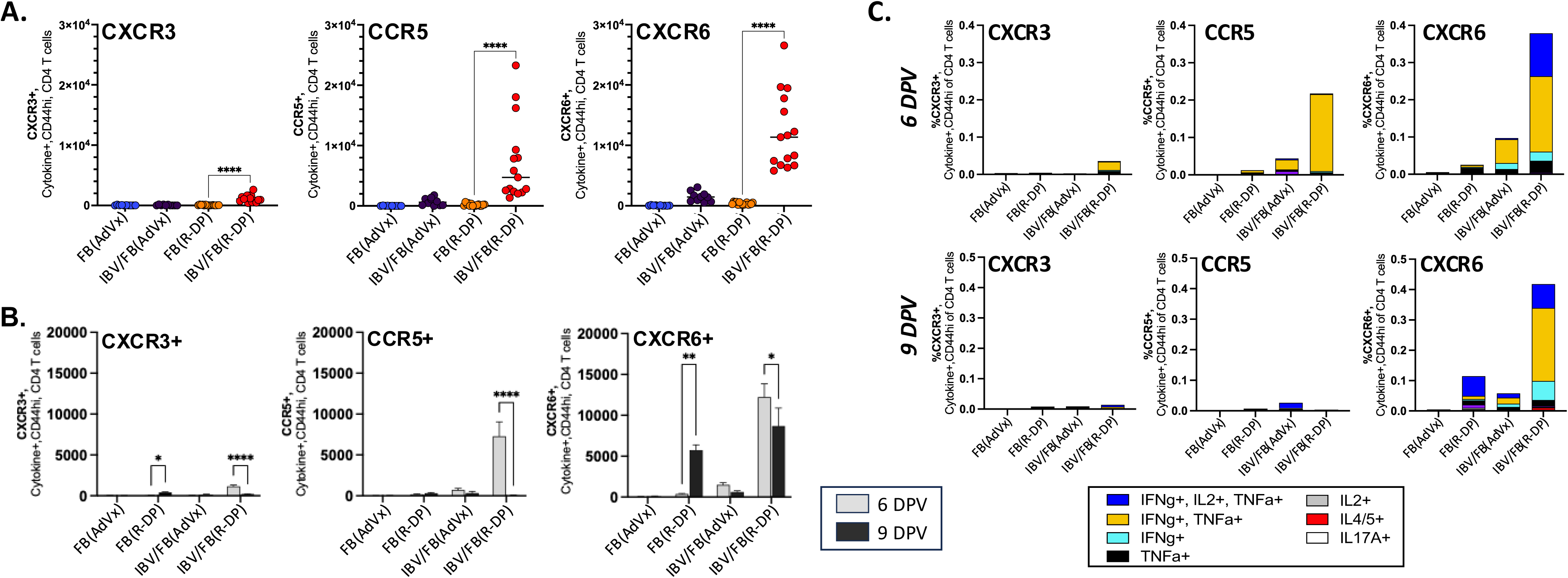
Chemokine receptor expression on HA-B specific CD4 T cells is enhanced in IBV memory mice. **(A)** After stimulation with HA-B peptide, total HA-B specific reactivity was determined by summing all cytokine producing CD4 T cells and fractional responses expressing the indicated chemokine receptor evaluated for the expression of CXCR3, CXCR6 or CCR5. Statistical significance was determined using One-Way ANOVA with Fisher’s LSD test. **(B)** CD4 T cells were purified and evaluated, and a direct comparison of chemokine receptor expression in HA B-reactive CD4 T cells at day 6 or day 9 days post vaccination is shown. Statistical significance was determined using two-Way ANOVA with Fisher’s LSD test. Bars reflect the mean with SEM. **(C)** The functional composition of the indicated chemokine receptor-expressing population is represented as stacked bars with each color representing a unique cytokine-producing population. The height of the bar reflects the mean of the fraction of the total CD4 T cell response that is chemokine receptor positive and produces the indicated cytokine after HA-B stimulation at 6dpv (top), compared to 9dpv (bottom).

The altered chemokine receptor expression profile early after vaccination supports the possibility that the HA-B specific CD4 T cells elicited by vaccination in previously infected host may egress in response to migratory cues between day 6 and day 9 post infection. If this is the case, we would expect the decline in overall response magnitude would manifest as a selective decline in the CD4 T cells expressing these receptors. To address this, we quantified cytokine-producing, chemokine-receptor expressing populations at 6 and 9 dpv (**Figure 4B**). A significant decline in the frequency of CCR5+, and to a lesser extent CXCR3+, cytokine producers in IBV/Flublok (RDP) mice was observed. We next evaluated the effector phenotype of these subsets of cells based on cytokine production to determine whether chemokine receptor expression is restricted to distinct subsets of CD4 T cells. In **Figure 4C**, the total frequency of cytokine producing, chemokine receptor expressing CD4 T cells at both timepoints was quantified to determine each subset’s contribution to the total chemokine receptor expressing populations of HA-B specific cells. This analysis revealed that chemokine receptor expressing HA-B specific CD4 T cells at day 6 post vaccination primarily consisted of the subset that produced both TNF-α and IFN-γ, depicted in gold. This was particularly striking in CCR5+, HA-B-reactive CD4 T cells. We speculated that the decline in response magnitude by 9 dpv in IBV memory mice, vaccinated with R-DP is due to the early recruitment of TNF-α, IFN-γ producing cells out of the vaccine-draining lymph node and into the periphery. Supporting this hypothesis is the almost exclusive loss of the CCR5 and CXCR3 expressing cells by day 9 (**Figure 4C**, bottom panel). It is interesting to note that the CXCR6 expressing cells did not decay. Most recent data suggest a major role of CXCR6 in recruitment of T cells to tumors ^32^, and studies in animal models suggest that vaccination of these hosts with tumor antigens facilitates tumor responses ^22,33^. It is possible that in the absence of tumor bearing host, these CD4 T cells do not rapidly migrate from the vaccine draining lymph node.

We considered two distinct mechanisms to explain the re-shaping of the CD4 T cell phenotype and fate in the host that has a history of infection. First, we speculated that infection-induced priming of CD4 T cells might “imprint” the elicited CD4 T cells for enrichment for cytokines such as IFN-γ and TNF-α, and the potential to express chemokine receptors associated with lung homing upon re-encounter with their cognate antigen in the vaccine-draining LN. The second but distinct possibility is that the infection-induced HA-B specific CD4 T cells have not been uniquely programmed for rapid upregulation of chemokine receptors and exit from the vaccine draining LN, but instead, when they are recruited to the vaccine-draining lymph node and become re-activated by encounter with vaccine bearing APC, the microenvironment of the lymph node is altered from that in naïve mice, perhaps with new mediators or innate cells inducing these changes in T cell phenotype and fate.

To distinguish between these possibilities, we examined responses to H3-derived epitopes, for which the CD4 T cells are naïve. There is no apparent cross reactivity between IBV elicited CD4 T cells and the dominant H3 peptide epitope studied here. In exploring the pattern of H3-specific CD4 T cell responses in naïve mice vs those mice with IBV infection induced memory, we predicted that if HA-B responses were uniquely associated with a change in phenotype and fate in a host with IBV-induced memory this would be consistent with infection-induced imprinting. In contrast, a concordance in the H3 and HA-B responses would support the model where the vaccine draining lymph node was modified by the recruitment of the infection-induced HA-B specific memory cells. Several of the hallmark changes observed in HA-B specific responses were compared with H3-specific responses. As shown in **Figure 5A**, H3-specific CD4 T cells elicited by Flublok administered with either immune activator displayed early responses at day 6, as had been observed with HA-B, with the magnitude of the response dictated by the adjuvant system. Also, when subsets of H3-specific CD4 T cells were compared for naïve mice and IBV memory mice, (**Figure 5B**), it was clear that the history of infection with IBV modified responses to H3. In the absence of memory, cytokine-producing cells increased from day 6 to day 9, while in the host with IBV infection history, CD4 T cells subsets specific for H3 were diminished by day 9. Particularly striking was the gain in CD4 T cells producing TNF-α and IFN-γ in response to Flublok/R-DP and the loss in IL-4 producing cells in the Flublok/AdVx vaccine responses to H3 in the hosts with IBV memory. The overall reshaping of the H3-specific CD4 T cells from the naïve host to the host with IBV memory is shown in **Figure 5C**, where, as had been done for HA-B responses, the subsets of cytokine-producing cells in the naïve mice compared to the host with IBV memory at the peak of each response is boxed. Although the overall response magnitude (indicated beneath each pie diagram) is lower for H3 than for HA-B, the qualitative shifts in response phenotypes are almost identical. In the host with an infection history, the H3 responses to Flublok/AdVx lose production of IL-4 and gain production of IFN-γ and TNF-α (**Figure 5C top**), while Flublok/R-DP become enriched for CD4 cells producing both IFN-γ and TNF-α (**Figure 5C bottom**).

**Figure 5.**
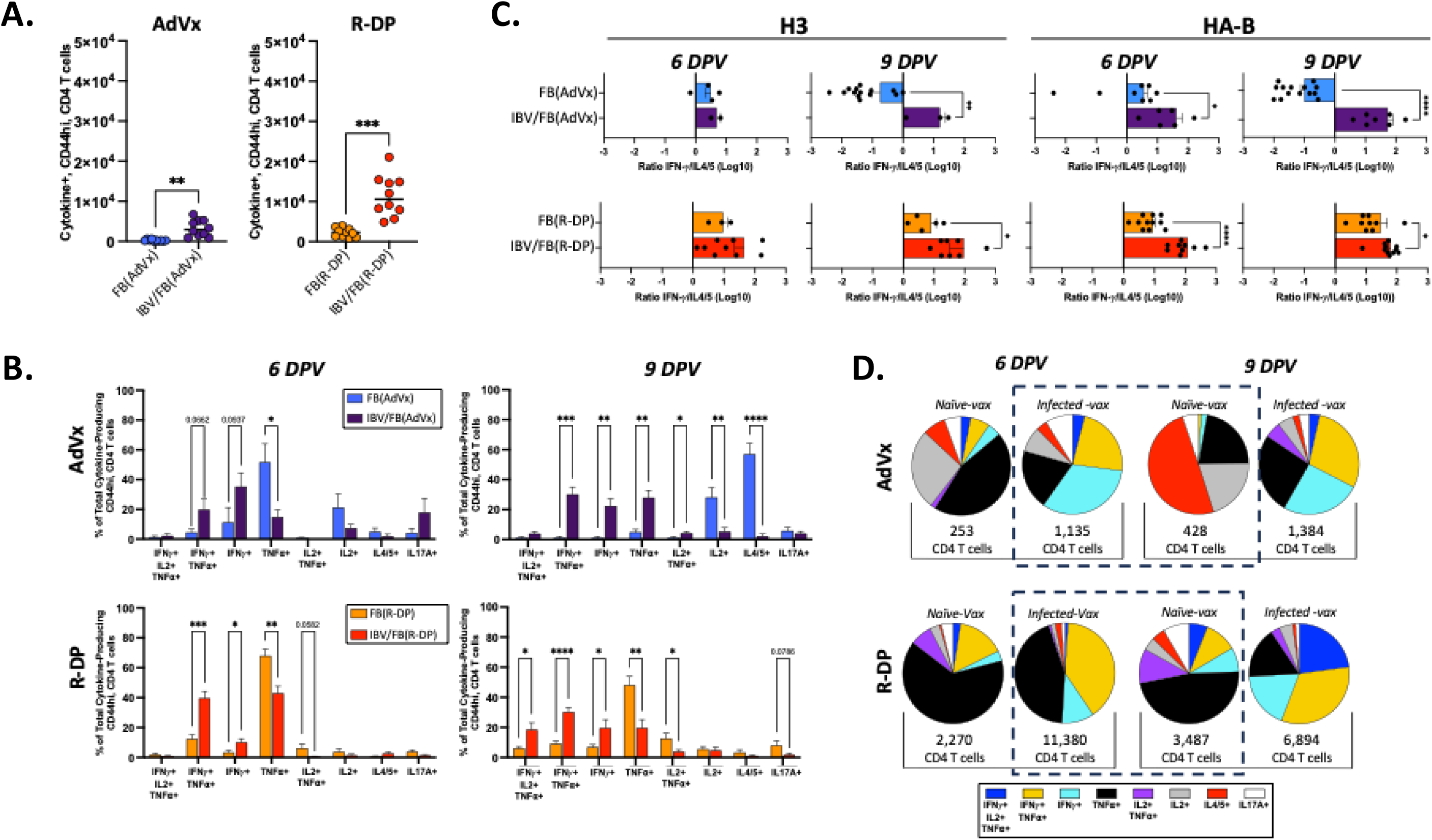
Remodeling of the Flublok elicited H3-specific CD4 T cells. Naïve mice or mice previously infected with IBV were vaccinated with Flublok, formulated with AdVx (IBV/Fb (AdVx), purple) or R-DP (IBV/Fb (RDP), red). At 6 days post-vaccination, H3 specific CD4 T cells were evaluated for cytokine production by ICS and flow cytometry. The total H3-reactive CD4 T cells, is shown, with statistical analysis using the unpaired t-test with Welch’s correction. **(B)** Fraction of the H3-specific CD4 T cell response for the cytokine-producing population at 6 (left) or 9 (right) days after vaccination with Flublok, formulated with AdVx (blue/purple, top) or R-DP (orange/red, bottom) in naïve mice and those previously IBV-infected. Statistics were performed by Two-way ANOVA with Fishers LSD test. **(C)** CD4 T cells from naïve mice or mice with IBV memory, vaccinated with Flublok were quantified for IFN-ψ or IL-4 production in response to HA-B (left) or H3 (right) stimulation. Ratios of IFN-ψ to IL-4/5 producers where IFN-ψ and IL-4/5 were identified by sequential gating are shown. Bars indicate the mean with SEM and statistical analysis was conducted using the Mann-Whitney, nonparametric t test. **D.** Pie graphs show functional composition of the H3-reactive CD4 T cell repertoire at 6 or 9 days after vaccination with Flublok (AdVx) (top) or Flublok (R-DP) (bottom) in naive mice or mice previously infected with IBV. The mean total of antigen-reactive CD4 T cells is noted below each pie.

The patterns of early (at day 6 post vaccination) chemokine receptor expression in H3-specific CD4 T cells shared similar early upregulation of CXCR3, CCR5 and CXCR6 in vaccinated hosts with infection history, which greatest in the hosts vaccinated with R-DP, but the overall magnitude of the responses is low, relative to HA-B, as shown earlier. **Figure 6A** and **Figure 6B**, shows that the kinetic patterns of accumulation and decay in H3-specific responses also mirrored HA-B specific CD4 T cells. Here, the peak of epitope-specific, chemokine receptor positive cells for both specificities occurred at day 6, dropping most sharply at day 9 post-vaccination in the previously infected and Flublok/R-DP vaccinated host for CXCR3 and CCR5 positive cells, with no impact on CXCR6 positive CD4 T cells.

**Figure 6.**
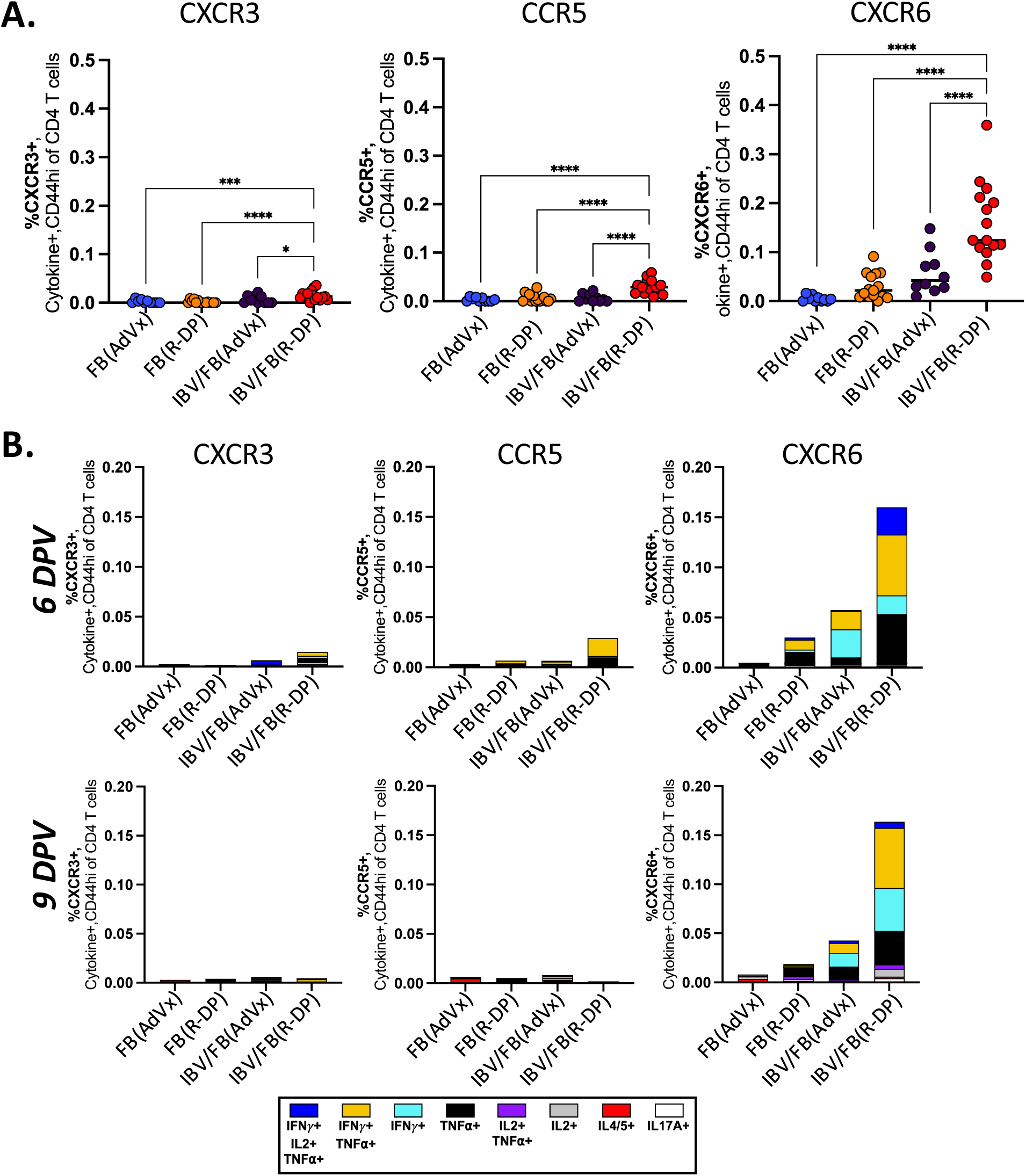
Vaccine-induced H3-specific CD4 T cells gain expression of chemokine receptors in host with history of IBV infection. **A.** The expression of the indicated chemokine receptors in H3-specific, cytokine-producing CD4 T cells is shown at day 6 post-vaccination. Naïve or previously IBV-infected mice were sampled after vaccination with Flublok, co-delivered with AdVx or R-DP. CD4 T cells, stimulated with H3 peptide were quantified by ICS and flow cytometry. The fraction of the H3–reactive CD4 T cell population expressing a given chemokine receptor, was determined after each unique cytokine producing population was evaluated for expression of CXCR3, CXCR6 or CCR5. These values were first back-calculated to total CD4 and then summed to determine the fraction of the total antigen-reactive population that was chemokine receptor positive. Statistical significance was determined using One-Way ANOVA with Fisher’s LSD test. The composition of each chemokine receptor-expressing population is represented in panel B as stacked bars with each color representing a unique cytokine-producing population. The height of the bar reflects the mean of the fraction of the total CD4 T cell response that expresses the indicated chemokine receptor and produces the cytokine after H3 peptide stimulation at day 6 or day 9 post-vaccination.

### In the context of memory, peripheral vaccination with Flublok (R-DP) leads to the significant accumulation of IFN-ψ producing HA-B specific CD4 T cells in the lung tissue

Peripheral vaccination has shown limited potential to generate CD4 T cells with lung trafficking potential. The chemokine receptor expression profiles of the HA-B and H3 specific CD4 T cells elicited by R-DP-Flublok in the context of immune memory to IBV led to decay of the cells that expressed lung homing chemokine receptors from the local dLN. To address the fate of the cells, vaccinated naïve and mice previously infected with IBV were sampled for HA-B specific CD4 T cells in lung tissues at 28 days. Cells in the lung vasculature were distinguished from cells in the lung tissue by IV labelling ^25^. Antigen specificity of the CD4 T cells was confirmed by quantifying and gating on the cells based on IFN-ψ cytokine production induced by HA-B peptide stimulation. An example of the gates utilized to identify CD45-, IFN-ψ populations for each cohort are shown **Figure 7A**. The results of these analyses, by frequency and total number have been quantified in (**Figure 7B top panel)**. Minor populations of HA-B specific CD4 T cells in the lung were detected with Flublok (R-DP) vaccinated naïve mice, but they were primarily within the vasculature and likely reflect circulating T cell populations. Compared to the magnitude of the response in IBV-infected mice at memory (IBV), we uncovered a significant, ∼5-fold increase in the frequency of HA-B reactive, IFN-*γ* producers in the lung tissue of IBV/Flublok (R-DP) mice. It is also important to note that infection memory alone is not sufficient to retain these populations of HA-B reactive CD4 T cells within the lung. The accumulation of HA-B specific CD4 T cells in the lung tissue at day 28 post vaccination at day 33 post infection was also observed in mice vaccinated at day 60 post infection (red hatched bars), arguing that residual antigen is unlikely to be necessary for lung retention of HA-B specific CD4 T cells. In agreement with this conclusion, is the finding that low but significant levels of H3-specific CD4 T cells are detectable at day 28 post vaccination after 31 days of memory (red solid bars) (**Figure 7B bottom panel).** The same trend is seen after a 60-day delay for vaccination (red hatched bars).

**Figure 7.**
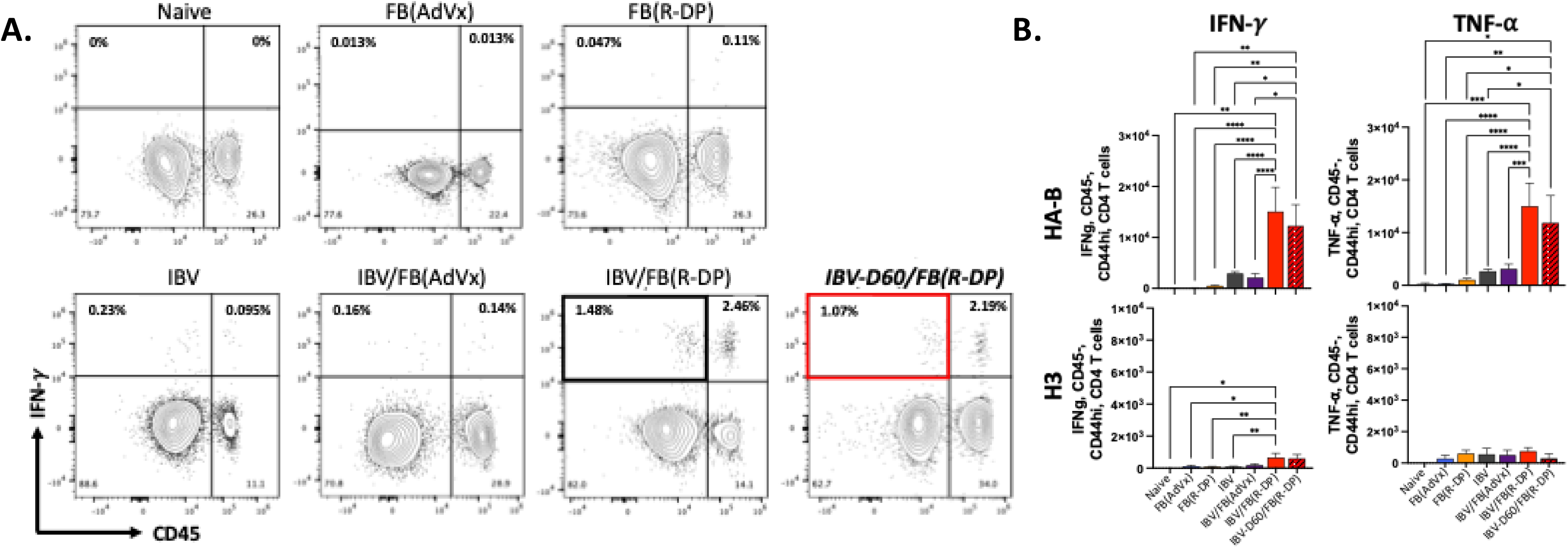
Tissue residency of vaccine elicited CD4 T cells in host with influenza infection vaccinated with Flublok-R-DP. To investigate the fate of vaccine elicited CD4 T cells, in naïve mice or mice previously infected with IBV, at 33 days post IBV infection, mice were vaccinated with the indicated Flublok formulation. Naïve mice were vaccinated in parallel. At 28 dpv, intravenous CD45 labeling was used to identify lung vasculature-localized cells. CD44hi, CD4 T cells were identified by sequential gating and the evaluated for IFN-ψ expression in the tissue (CD45-) or vasculature (CD45+), as demonstrated in the representative plots (Panel A). Similar gating strategy was used to identify or TNF-α or IL-4/5 producers. These analyses on individual mice are represented as bar graphs, comparing the total quantity of IFN-ψ, TNF-α and IL4/5 producers after stimulation with either HA B (top) or H3 (bottom). Shown are the numbers of HA-B (top) or H3-specific (bottom) CD4 T cells producing the indicated cytokines in the lung tissue at 28 days (open bars) or 60 days (hatched red bar) after infection. Bars indicate the mean with SEM and statistical analysis was conducted using One-way ANOVA with Tukey’s multiple comparison’s test.

The profound impact of immune memory from infection on CD4 T cell responses to both HA-B and H3 prompted us to examine the impact of serum antibody to HA. Shown in **Figure 8** are the results of ELISA to evaluate IgG responses to HA-B and H3 at day 28 post vaccination. In panel A, are the results of analyses of pooled sera from each of the 4 cohorts that varied by the pre-existing memory and adjuvant used for vaccination as well as the unvaccinated mice that had memory from IBV infection. Shown beneath these titration curves are the analyses of single mice where the dilution chosen was below the plateau values (1:1000 for HA-B and 1:100 for H3). These ELISA enabled statistical treatment of data. These data, particularly notable in the Flublok(AdVx) vaccinated mice demonstrate that not only is the HA-B specific antibody response to vaccination enhanced in hosts with memory from influenza B virus infection, for which there is both B cell and CD4 T cell memory, but that the same enhancement in antibody titers is seen for H3 specific antibody responses, for which there is no B cell or CD4 T cell memory. Collectively these data demonstrate that immune memory derived from influenza infection dramatically reshapes the adaptive CD4 T cell and B cell responses broadly, for both the antigens derived from the infecting virus and other antigens contained in the multivalent vaccine.

**Figure 8.**
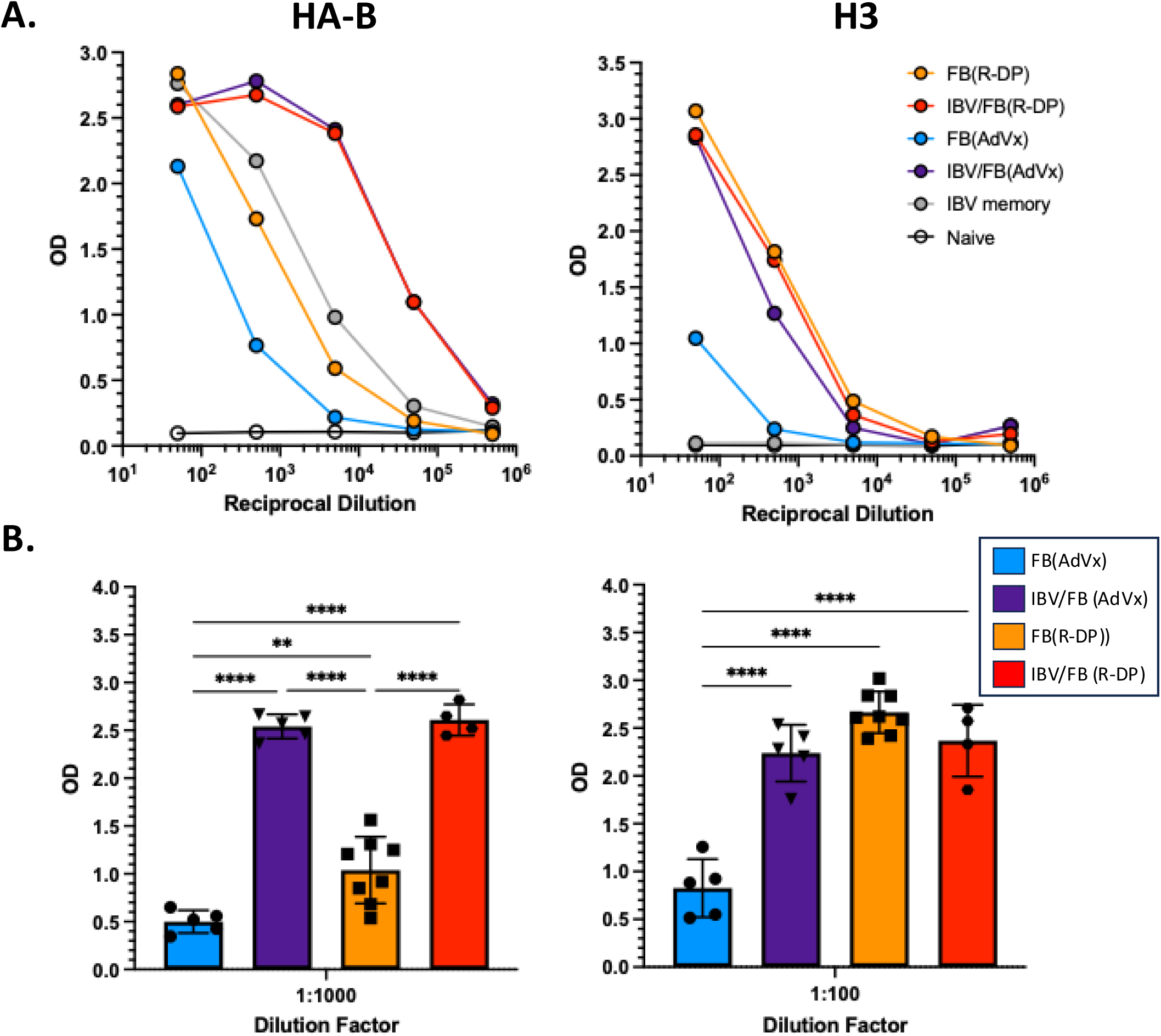
HA-specific serum IgG responses at day 28 post-vaccination in naïve mice or in mice with infection history. Naïve mice or mice with IBV infection history were immunized with Flublok formulated with either R-DP or AdVx. Sera collected at day 28 post-immunization were tested by ELISA for HA-specific IgG. (**A**) Dilution curves from pooled sera from >5 mice for each group for HA-B (left) and H3 (right), including vaccination only and infection plus vaccination conditions and IBV memory. OD values were measured at 405 nm across serial dilutions starting at 1:50 in 10-fold increments, plotted on a log scale. **Panel B** shows bar graphs representing the OD of ELISA from individual mice at dilutions on the linear segment of the dilution curve shown in A (1:1000 for HA-B, left and 1:100 for H3, right). Each point represents an individual mouse, with error bars indicating standard deviation. Group are indicated as FB(AdVx) (blue, n = 5), IBV/FB(AdVx) (purple, n = 5), FB(R-DP) (orange, n = 8), and IBV/FB(R-DP) (orange, n = 4). Data are represented as mean response with SD and one-way ANOVA with multiple comparisons. Statistically significant differences are denoted (****p < 0.0001).

## Discussion

In this study, to better model human immunity to influenza vaccination, we have addressed the impact of immune memory from infection on vaccine responses to a licensed recombinant vaccine administered with candidate immunomodulatory systems. These studies revealed striking differences in CD4 T cell response kinetics, magnitude, phenotype and fate post-vaccination in the host with infection-induced memory. The response peaks early in the vaccine draining lymph node by day 6 post vaccination and thereafter decays, while in the naïve mouse, CD4 T cells continue to accumulate until day 9. Most strikingly, when administered with the nanolipoprotein R-DP, the host with immune memory elicits a CD4 T cell response dominated by TNF-α− and IFN-ψ-producing CD4 T cells that express the chemokine receptors CXCR6, CCR5 and CXCR3. The TNF-α and IFN-ψ producing cells expressing either CXCR3 or CCR5 rapidly decay in the vaccine dLN between day 6 and day 9 post vaccination, and this decline is kinetically linked to the appearance of epitope specific IFN-ψ /TNF-α producing cells in the lung tissue that persist for at least 30 days post vaccination. The simplest model to explain this decay is that the chemokine receptor positive cells leave the vaccine draining lymph node and some fraction home to the lung tissue where they establish residency. However, we have not traced this directly, and therefore cannot exclude that some of these cells, distinguished by expression of these receptors, die after the acute expansion. Remarkably, this reshaping of the CD4 T cell response to vaccination in the host with infection history does not require priming by infection. CD4 T cells from an alternate HA (H3) contained in the vaccine distinct from the virus used for infection (IBV) undergoes the same alteration in kinetics, cytokine patterns and cell surface phenotype in the response. Likely as a consequence of this reshaping of the H3 specific CD4 T cell response is an enhanced B cell response to H3. We have no evidence that the H3 specific responses we detect are due to cross reactivity of the CD4 T cells elicited by IBV infection with the dominant H3 peptide used in these assays.

From these data, we speculate that the most likely mechanism to explain this dramatic reshaping of the CD4 T cell vaccine response is an alteration in the microenvironment in the vaccine draining lymph node upon recruitment of memory cells from the IBV infection. Although we have only tracked CD4 T cell responses, it is possible that other cells recruited to the lymph node within the memory containing host contribute to the presumed alteration in the microenvironment. These cells could include the infection primed CD8 T cells or B cells or recruitment of alternate APC or innate cells ^34^. These issues are the topic of current and future investigations.

However, of relevance to consider now are several issues. First, will the vaccine elicited CD4 T cells that home to the lung persist there and serve as sentinel cells for future infections? There is much interest in developing intranasal vaccines to generate mucosal immunity in the respiratory tract ^35–37^ but these are not without risk or concerns ^38,39^. If a peripherally administered protein-based vaccine, administered with a safe immunomodulatory reagent, could foster long lived protective CD4 T cells in the respiratory tract, this could have considerable safety and efficacy advantages. The key features of the R-DP immunomodulatory agent identified thus far is an early type I IFN responses initiated by engagement of TLR 7/8. The ideal type of formulation would potentiate many arms of the immune responses, some studied here, include rapid help for antibody responses, direct effector function via cytokines with known antiviral effects and potentiation of lung homing.

Another important issue to consider based on our findings here are whether and how the consequences of single infection history studied here on vaccine responses will be modified by the repeated and diverse confrontation with influenza antigens via both infection and vaccination that takes place in humans. This type of complex immune memory is difficult to establish in small animal models with limited longevity. However, because R-DP has shown an excellent safety profile in humans, it may be possible to study the multiple features of vaccine-induced immunity in adults with this cationic lipid nanoparticle relative to the comparators studied here.

## Supporting information

Main Manuscript text and figure

## Acknowledgements

This work was supported with Federal funds from the National Institute of Allergy and Infectious Diseases, National Institutes of Health, Department of Health and Human Services, NIAID Collaborative Influenza Vaccine Innovation Centers (CIVIC – CIVR-HRP) Contract number 75N93019C00052, U01 AI14461602S1 and Centers of Excellence for Influenza Research and Response (CEIRR-CIDER) contract number 75N93021C00018 sub-contracts to AJS. This work was supported by Federal Funds from the National Institutes of Health Pathogenesis T32 5T32AI118689-07 and Immunology T32 5T32AI7285-34 to CLW.

## Author Contributions

Conceptualization, AJS; methodology, CLW, LM, SKG; formal analysis, CLW, AJS; investigation, CLW, AJS; resources, SKG; data curation, CLW, KAR, AJS; writing—original draft preparation, AJS, CLW; writing—review and editing, AJS, CLW, KAR; visualization, AJS, CLW, KAR; supervision, AJS; funding acquisition, AJS. All authors have read and agreed to the published version of the manuscript.

## Data Availability

The full complement of data accumulated for these studies is available upon request.

