## Supplementary material for "Re-shaping the immune response to influenza vaccination in a host with immune memory from influenza": Main Manuscript text and figure

Supplementary Data

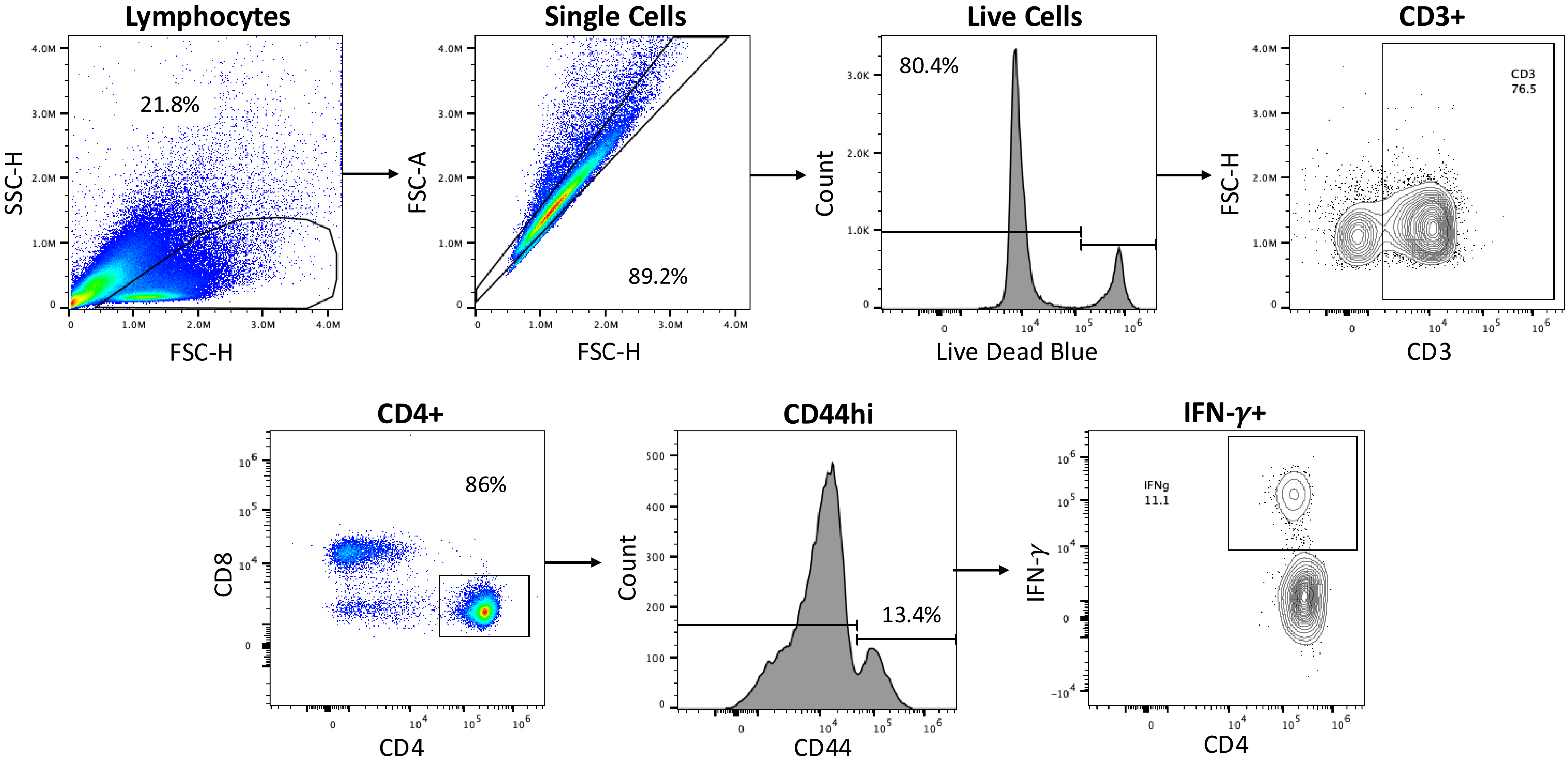


**Supplemental Figure 1**. Representative flow plots for gating strategy utilized to identify antigen-reactive CD4 T cells

| Antibody | Fluorophore | Clone | Company | Catalog # |
|---|---|---|---|---|
| IL-17a | BUV395 | TC11-18H10 | BD | 565246 |
| Live Dead | Live Dead Blue | | Invitrogen | L23105 |
| B220 | BUV496 | RA3-6B2 | BD | 612950 |
| CD11a | BUV563 | 2D7 | BD | 741225 |
| CD49d | BUV615 | R1-2 | BD | 752352 |
| CCR7 | BUV661 | 4B12 | BD | 741677 |
| CD69 | BUV737 | H1.2F3 | BD | 612793 |
| CD3 | BUV805 | 17A2 | BD | 569192 |
| CXCR6 | BV421 | SA051D1 | BioLegend | 151109 |
| CD62L | BV510 | MEL-14 | BioLegend | 104441 |
| CCR5 | BV605 | C34-3448 | BD | 743697 |
| TNFa | BV650 | MP6-XT22 | BD | 563943 |
| CD49a | BV711 | Ra31/8 | BD | 564863 |
| IL-2 | BV750 | JES6-5H4 | BD | 566363 |
| IFN-$\gamma$ | BV786 | XMG1.2 | BD | 563773 |
| CD8a | FITC | 53-6.7 | BD | 553031 |
| CXCR3 | PE-eFluor 610 | CXCR3-173 | Invitrogen | 61-1831-82 |
| CD107a | PerCP-eFluor 710 | 1D4B | Invitrogen | 46-1071-82 |
| IL-4/IL-5 | PE | 11B11 | BioLegend | 504104 |
| | | TRFK5 | BioLegend | 504304 |
| CD4 | PE-Cy5 | RM4-5 | BD | 553050 |
| CD103 | PE-Cy7 | M290 | BD | 567593 |
| CD45 | APC | 30-F11 | TONBO | 20-0451-U100 |
| CD44 | APC-Cy7 | IM7 | TONBO | 25-0441-U100 |

Supplemental Table 1. List of antibodies used in the study.
